# An Observational Study of the Impact of Systemic B-cell Depletion on Cervicovaginal Mucosal Environment

**DOI:** 10.64898/2026.04.16.718227

**Authors:** Ofri Bar, Meena Murthy, Katherine Cosgrove, Yusra Saidi, Wafae El-Arar, Miles Goldenberg, Gabriel Sauvage, Agnes Bergerat, Briah Cooley Demidkina, Karen Laliberte, Jiawu Xu, Grace Pierson, Douglas S Kwon, John Niles, Moran Yassour, Caroline M. Mitchell

**Affiliations:** Vincent Center for Reproductive Biology, Massachusetts General Hospital, Boston, MA; Department of Microbiology and Molecular Genetics, Faculty of Medicine, Hebrew University of Jerusalem; Harvard Medical School, Boston, MA; Vasculitis and Glomerulonephritis Center, Massachusetts General Hospital, Boston, MA; Ragon Institute of MGH, MIT, and Harvard, Cambridge, MA; Division of Infectious Diseases, Massachusetts General Hospital, Boston, MA; Division of Nephrology, Massachusetts General Hospital, Boston MA; The Rachel and Selim Benin School of Computer Science and Engineering, Hebrew University of Jerusalem; Department of Obstetrics and Gynecology, Mass General Brigham, Boston MA

## Abstract

**Importance:** Emerging data show that B-cell depleting chemotherapies, which are increasingly used to treat autoimmune disorders and multiple sclerosis, can be associated with mucosal side effects such as inflammatory vaginitis.

**Objective:** Evaluate the impact of rituximab treatment on vaginal mucosal immune markers, endocervical immune cell populations and vaginal microbiome.

**Design:** Cross-sectional observational study conducted between 2022 – 2024.

**Setting:** Academic medical center, Boston Massachusetts.

**Participants:** We enrolled women aged >18 years who were either 1) receiving rituximab for autoimmune renal disease or were 2) healthy controls

**Exposure:** Treatment with rituximab, an anti CD20 monoclonal antibody.

**Main outcome and measure:** We compared endocervical immune cell populations, vaginal fluid immune markers, vaginal fluid immunoglobulins and vaginal microbiome composition between individuals being treated with rituximab and healthy controls.

**Results:** We enrolled 26 women treated with rituximab for autoimmune renal disease and 26 healthy controls. Median circulating and endocervical B-cell and plasma cell proportions were significantly lower in treated participants compared to controls. Median vaginal fluid IgA concentrations were significantly lower in participants treated with rituximab, while ILE, IgM, IgG1, IgG2, IgG3 and IgG4 were not different between groups. Total T cell frequencies were similar between groups, but the proportion of activated T cells (CD4+CD38+HLADR+) was significantly lower in people treated with rituximab. Concentrations of IL10, IL13, IL17, IL21, IL23, IL4, ITAC and TNFa were elevated in vaginal fluid from the rituximab group, while IL-8 was lower. A CST-IV-C, low-*Lactobacillus* pattern of vaginal microbiota was more common in the rituximab group.

**Conclusions and Relevance:** Systemic B-cell depletion is associated with reduced vaginal fluid IgA, a more diverse microbiome composition, and increases in many vaginal fluid immune markers compared to healthy controls. The reduction in vaginal fluid IgA may provide opportunities for vaginal bacteria to induce inflammation.

**Key points:** *Question:* How does circulating B-cell depletion impact the vaginal microenvironment?

*Findings:* In this cross-sectional study of 52 women, B cell and plasma cell proportions were significantly lower in both blood and vaginal mucosa among rituximab-treated participants compared to healthy controls. Vaginal IgA concentrations, but not other immunoglobulins, were significantly lower in rituximab treated participants. In treated participants, vaginal cytokine concentrations were elevated, and microbiome composition shifted toward non–*Lactobacillus*-dominant communities. In six people with inflammatory vaginitis, both circulating and endocervical B cells were lowest in people with the most severe symptoms.

*Meaning:* Systemic B cell depletion is associated with alterations in vaginal mucosal immune markers and microbiome composition which increase local inflammation.

## Introduction

Rituximab, a chimeric monoclonal antibody targeting CD20 is widely used to treat autoimmune diseases by depleting B lymphocytes. B cell depletion effectively reduces disease activity in conditions like rheumatoid arthritis,^1^ lupus,^2^ and multiple sclerosis^3^. Autoimmune diseases disproportionately affect women; thus it is important to understand the impact of B-cell depleting medications on reproductive health. There are limited data on how rituximab impacts host immune function in the female genital tract (FGT).

Mucocutaneous complications of the vagina and vulva have been described in women being treated with rituximab for cancer^4^. We previously reported the development of inflammatory vaginitis among patients on long-term rituximab treatment for autoimmune disorders, characterized by unexplained vaginal symptoms encompassing vaginal discharge, pain, irritation, and dyspareunia^5^. Among people in our cohort who stopped rituximab and had return of systemic circulating B lymphocytes, most reported resolution of symptoms in contrast to those who remained on treatment. Since then, two additional publications have described inflammatory vaginitis in women being treated for multiple sclerosis with B-cell depleting chemotherapy, and a similar resolution was seen with cession of treatment and B-cell restoration^6,7^.

B cells in the female genital tract are an understudied cell type, and little is known about their role in regulating mucosal immunity and host defenses at this surface. In this study, we investigated the impact of systemic B-cell depletion on the cervicovaginal mucosal immune environment and the vaginal microbiome by comparing patients receiving rituximab to healthy controls.

## Methods

### Study design

This study was approved by the ethics committee of the Massachusetts General Brigham Institutional Review Board (2021P001522 and 2023P000472). For the rituximab study group, patients with an autoimmune disorder impacting their kidneys receiving rituximab infusions at the Massachusetts General Hospital vascular glomerulonephritis clinic were offered enrollment between July 2021 and April 2023. Healthy controls were recruited from the community through flyers and research advertisements. Eligible participants were women over the age of 18. Exclusion criteria for healthy controls included: pregnancy, vulvar dermatosis, regular use of antibiotics (i.e. daily or post-coital prophylaxis for UTI), recent probiotic use (< 30d), abnormal pap within the past 1 year. Women who had used probiotics in the past 30 days were offered enrollment but rescheduled for a time > 30 days after last use. All participants attended at least a single study visit, and a subset of participants in the rituximab group attended two study visits over a six-month period. During each visit, participants completed surveys about medical and gynecologic history, recent sexual activity and vaginal hygiene practices, and demographic information. Participants were asked to rate the severity of symptoms of vulvar itch/burn/irritation, vaginal itch/burn/irritation, vulvar pain, vaginal pain, vaginal discharge, and vaginal dryness from 0 (none) to 5 (severe).

### Sample collection and processing

Participants provided vaginal swabs and placed a disposable menstrual cup (Softdisc, Flex Company) in the vagina for 20 min to collect vaginal fluid. The menstrual cup was placed in a sterile 50 ml conical tube and transported to the laboratory on ice. The sample was weighed, and the weight of a clean tube and menstrual cup subtracted to get specimen weight. The tube was centrifuged at 810xg at 4 °C for 10 min and then the menstrual cup removed with sterile forceps, scraping any adherent secretions into the tube with a sterile spatula. Sterile saline (500uL) was added and the vaginal fluid was homogenized by aspirating it through a blunt 16-gauge needle 20 times, then aliquoted and stored at -80 °C.

### Vaginal fluid gram stain

A vaginal swab was smeared on a glass slide, which was allowed to dry and then was heat fixed and Gram stained. Bright field microscopy was used to assess the ratio of white blood cells (WBC) to epithelial cells in 400X fields, and each slide was scored as no WBC, < 1 WBC/epithelial cell, or > 1 WBC/epithelial cell.

### Immune cell analysis

An endocervical cytobrush was collected by placing the brush in the cervical os and rotating twice. The sample was placed in R10 media (500 mL RPMI media + 5mL 200 mM L-glutamine + 5mL 1 M Hepes + 25000 IU Pencillin/25000 mg Streptomycin + 10% fetal calf serum) at 4 °C and processed within 2 h for flow cytometry. Cells were stained with LIVE/DEAD fixable blue dead cell stain dye and fluorescent monoclonal antibodies specific for the following human surface markers: CD45 (Alexa700), CD8 (APC-H7), CD56 (APC), CD3 (PE-CF594), CD66b (PE), CD4 (BV786), CD14 (BV605), CD19 (BV510), CCR5 (BV421), CD11c (BUV737),CD16 (BUV395) and the activation markers HLA-DR (FITC) and CD38 (BV650)^8^. The samples were then fixed with 1% paraformaldehyde (PFA) and analyzed on a flow cytometer within 48 h.

Flow cytometry was performed on the 4-laser LSR2 (LIVE samples) (4 L LSR2), 4-laser LSR2 (4R LSR2), and the 5-laser LSR Fortessa (5 L Fortessa) (BD). We used the following controls for each experiment: (1) an unstained sample, (2) a fluorescence minus one control for a subset of markers and (3) calibration (“rainbow”) beads. These allowed us to ensure comparability between experiments. Data analysis was conducted using FlowJo Version 7.0.0 (FlowJo Enterprise). We defined B cells as CD45+CD3-CD19+, and plasma cells as CD45+CD3-CD19+CD38+/CD138+^9^. The gating diagram is displayed in Supplemental Figure 1.

### Cytokine analysis

To assess changes in the concentration of vaginal mucosal cytokines, we used quantitative fluorescent flow cytometry (Luminex®). Processed frozen vaginal fluid samples from the disposable menstrual cup were thawed on wet ice, and 100 uL of each sample was transferred to a 96-well plate. The samples were then diluted with 100 uL of PBS and centrifuged at 1000xg for 15 min. Afterwards, the supernatant from each well was transferred to a Millipore® 0.22 μm MultiScreenHTS GV Filter Plate that was centrifuged at 2451 x g for 1 h. Cytokine quantities in the filtered supernatant samples were measured with a MILLIPLEX® MAP Custom 20-Plex High Sensitivity Kit on a FlexMap 3D Luminex machine with xPONENT software. 20 differents analytes were measured in the kit: ITAC, IFN-γ, IL-10, MIP-3α, IL-12 (p70), IL-13, IL-17, IL-1α, IL-1β, MIG, IL-21, IL-4, IL-23, IL-5, IL-6, IL-8, IP-10, MIP-1α, MIP-1β, and TNF-α. Kit reagents were diluted by a factor of 1/3, and all other steps were done according to the manufacturer’s protocol. Samples that had analytes with bead counts below 20 were re-run. During analysis the outcome data was normalized for the menstrual cup volume. Values below the lower limit of detection in the assay were recorded as half of the lowest standard concentration for that analyte. Similarly, concentrations above the detectable limit were recorded as 2x the highest standard concentration.

### Immunoglobulin analysis

Immunoglobulin quantities in the filtered supernatant samples were measured with a MILLIPLEX® MAP Custom 7-Plex High Sensitivity Kit on a FlexMap 3D Luminex machine with xPONENT software. Softcup fluid was pre-processed as for cytokine analyses. The immunoglobulins measured included: IgA, IgM, IgE, IgG1, IgG2, IgG3 and IgG4. Samples that had analytes with bead counts below 20 were re-run. Samples with values above the upper limit of detection were further diluted and re-run. During analysis the outcome data was normalized for the menstrual cup volume. Values below the lower limit of detection in the assay were recorded as half of the lowest standard concentration for that analyte.

### Microbiome analysis

Vaginal swabs were eluted in 400uL sterile saline, centrifuged and the pellet submitted for DNA extraction. DNA underwent 16S rRNA gene sequencing using Illumina MiSeq of the V4 region of the 16S rRNA gene, and taxonomic assignment was performed as previously described^10,11^. In stacked bar plots, only the 20 most prevalent taxa are represented. The microbial compositional analysis was performed using R (Version 4.2.2). Taxonomy data was aggregated at the genus level and low abundance taxa (< 0.5% prevalence) were filtered out. Samples were assigned to a microbial Community State Type (CST) using *VAginaL community state typE Nearest CentroId clAssifier* - VALENCIA^13^.

### Statistical analysis

Our sample size was determined by the number of women receiving rituximab in the MGH Vasculitis and Glomerulonephritis clinic who agreed to participate during the study period. We enrolled healthy controls in a 1:1 ratio with our cases but did not match on any specific criteria. Metadata were compared between participants receiving rituximab treatment and healthy controls using chi square, t-test or Mann-Whitney U test as appropriate.

Alpha diversity was assessed using the Shannon Diversity Index and was calculated using the Diversity function which is part of the Vegan package. Symptom severity was grouped into categories: None (0), mild (1), and moderate to severe (2). We then created a composite symptom score (range 0-6) by counting the number of symptoms rated moderate-severe.

Inflammatory vaginitis was defined as having moderate to severe vaginal discharge and either moderate to severe vaginal pain or >1 WBC/epithelial cell in the vaginal fluid Gram stain assessment.

When comparing immune cell proportions, cytokines, or immunoglobulins between rituximab and control groups, we included only data from the first timepoint for the rituximab group. When evaluating the association between microbiome and study group all samples were used. We compared the proportions of community state types (CST) between groups using chi square and adjusted for menopause using logistic regression. Statistical analyses were performed using either Stata or R, with a p-value of less than 0.05 considered statistically significant.

Associations between time since last rituximab dose, total number of rituximab doses and vaginal fluid IgA concentration were examined using loess regression.

## Results

We enrolled 26 individuals receiving rituximab and 26 healthy controls in the study. Among those receiving rituximab, 19 provided samples at two different visits. Of the people who did not provide a second sample, four declined a second visit, one became pregnant, and two were unable to complete a second visit before study closure. For the cross-sectional analysis, we included only one sample per participant. For rituximab-treated participants we selected the first visit sample except for four participants who were initiating treatment at their first visit, for whom the second-visit sample was used(Table 1). Out of 26 individuals receiving rituximab, a total of 6 (23.1%) were identified as having inflammatory vaginitis. Of these, 5 participants fulfilled the study definition (all had moderate-to-severe discharge with >1 WBC, and 3 also reported vulvovaginal pain), while 1 participant entered with a prior clinical diagnosis.

**Table 1.**
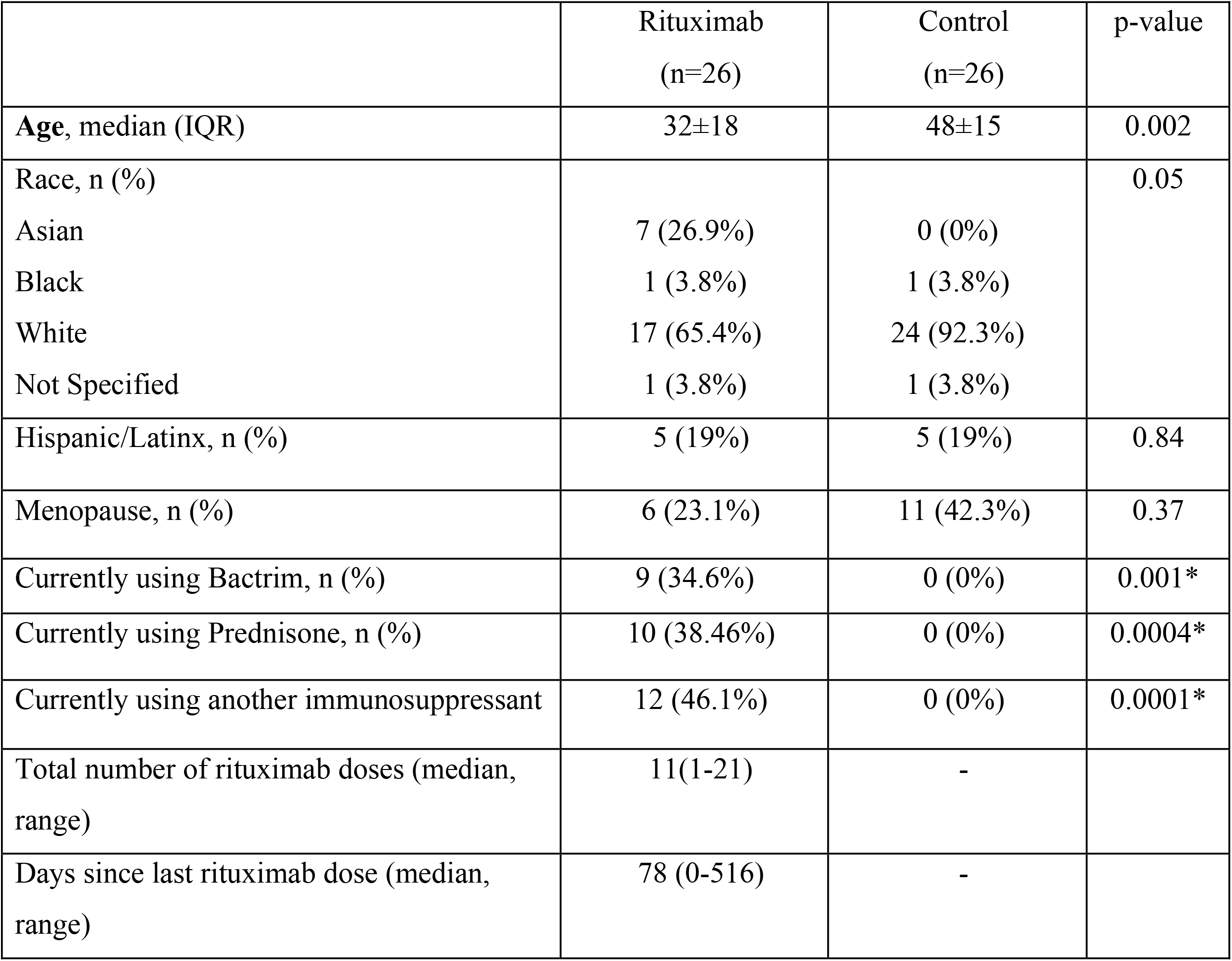
Patient demographics and use of medication at the time of enrollment.

|  | Rituximab<br>(n=26) | Control<br>(n=26) | p-value |
| --- | --- | --- | --- |
| **Age**, median (IQR) | 32±18 | 48±15 | 0.002 |
| Race, n (%) |  |  | 0.05 |
| Asian | 7 (26.9%) | 0 (0%) |  |
| Black | 1 (3.8%) | 1 (3.8%) |  |
| White | 17 (65.4%) | 24 (92.3%) |  |
| Not Specified | 1 (3.8%) | 1 (3.8%) |  |
| Hispanic/Latinx, n (%) | 5 (19%) | 5 (19%) | 0.84 |
| Menopause, n (%) | 6 (23.1%) | 11 (42.3%) | 0.37 |
| Currently using Bactrim, n (%) | 9 (34.6%) | 0 (0%) | 0.001* |
| Currently using Prednisone, n (%) | 10 (38.46%) | 0 (0%) | 0.0004* |
| Currently using another immunosuppressant | 12 (46.1%) | 0 (0%) | 0.0001* |
| Total number of rituximab doses (median, range) | 11(1-21) | - |  |
| Days since last rituximab dose (median, range) | 78 (0-516) | - |  |

We compared the proportions of immune cell populations between control participants and patients treated with rituximab in the periphery and in the endocervical mucosa (Figure 1). As expected, the proportions of B cells (CD45+CD3-CD19+) and plasma cells (CD19+ CD38+/ CD138+) were significantly lower in both the periphery and the endocervix of patients treated with rituximab compared to controls (Figure 1A, 1B). The difference in B-cell proportion between the treated vs. control group was smaller in the endocervix than the periphery, though the comparison did not reach statistical significance (Figure 1A). This was also true for plasma cells (Figure 1B), but proportions of neutrophils (CD66b+) showed similar differences between treated vs. controls participants in the periphery and in the vaginal mucosa (Figure 1C).

**Figure 1.**
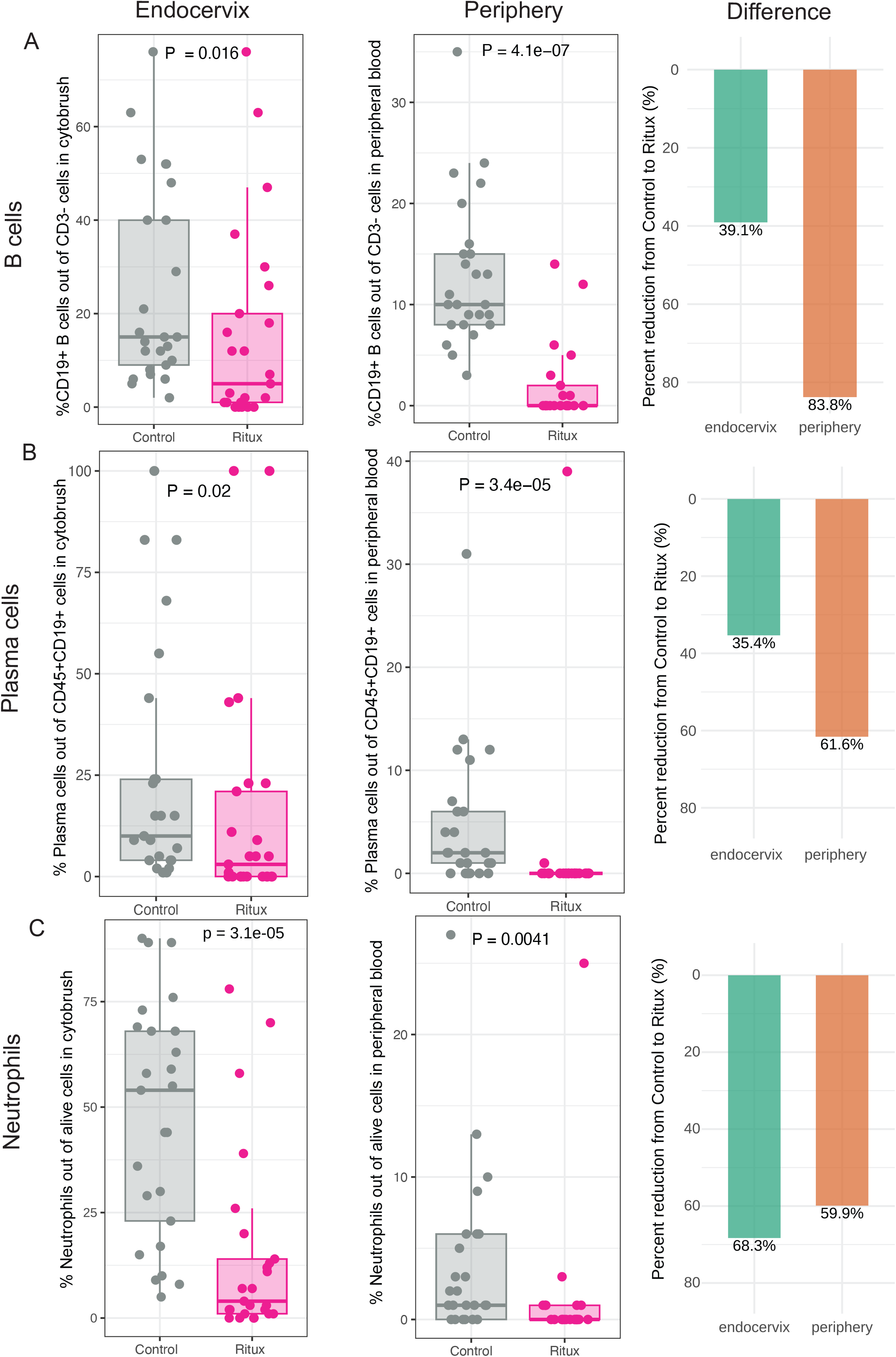
Immune Cell Proportions in control vs. rituximab-treated participants. Comparison of median proportions of B cells (A) in endocervical (left panel) and peripheral blood (middle panel) samples demonstrates a lower proportion in rituximab-treated vs. control participants for both compartments. However, there is a smaller difference (right panel) between control vs. rituximab treated participants in the endocervix than in the periphery. This difference between the compartments was less pronounced for plasma cells (B), and neutrophils (C), both of which were significantly lower in rituximab-treated participants. Error bars represent 25th and 75th percentiles. This analysis included 25 control and 25 rituximab-treated participants.

Proportions of CD45+CD3+CD4+ and CD45+CD3+CD8+ endocervical T cell populations were not different between patients receiving Rituximab and controls (Supplemental figure 2).

However both endocervical CD4+ CD38+ HLA-DR+ and CD4+ CD38-HLA-DR+ activated T cell populations were significantly lower in participants receiving rituximab compared to controls (Supplemental Figure 3A). Endocervical CD8+CD38+ HLA-DR+ and CD8+CD38-HLA-DR+ populations were also significantly lower in participants receiving rituximab compared to controls (Supplemental Figure 3B). These findings suggest that in patients receiving rituximab, the proportion of total T cells population is preserved, but the balance between activated and non-activated T cells is altered compared to healthy controls.

Vaginal fluid IgA concentrations were significantly lower in participants receiving rituximab treatment compared to healthy controls, while no significant differences between groups were observed for other immunoglobulin subtypes (Figure 2A-G). Of the 20 cytokines and chemokines measured we found vaginal fluid IL-10, IL-13, IL-17, IL-4, IL-23, ITAC and TNF-α were all significantly higher in people receiving rituximab vs. controls (Supplemental Figure 4). Only IL-8 concentrations were significantly higher in the control group compared to people receiving rituximab (Supplemental Figure 4). People receiving rituximab were more likely to have CST-IV-C (non-*Lactobacillus* dominant, diverse) communities, even after adjusting for menopause (Supplemental Figure 5). Of note, 9/26 (34%) of rituximab-treated participants were also taking Bactrim for prophylaxis (Table 1).

**Figure 2.**
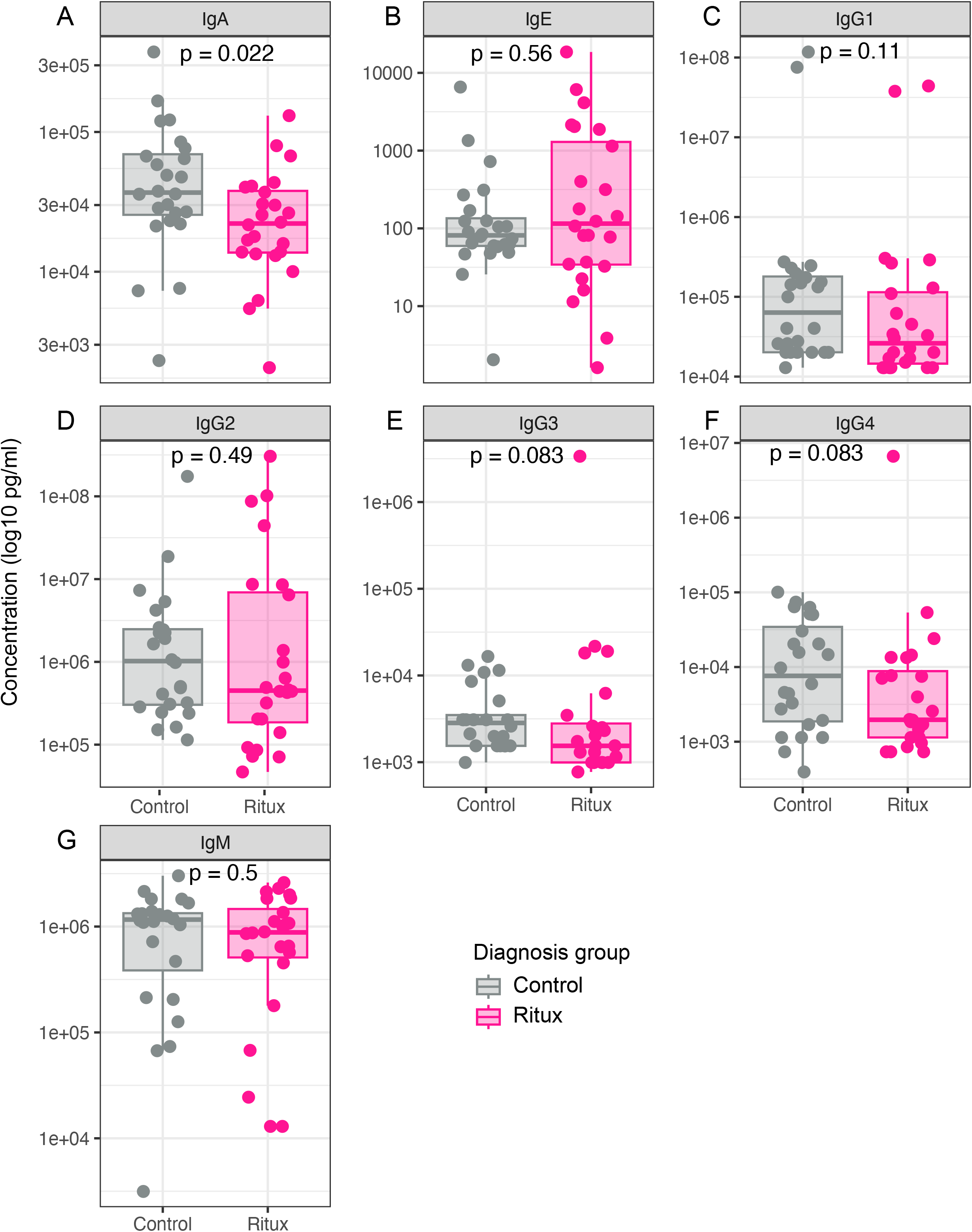
Vaginal fluid immunoglobulin concentrations. In vaginal fluid, concentrations of IgA (A) were significantly lower in rituximab-treated participants vs. control participants, while no difference was seen in IgE (B), IgG subclasses (C-F) or IgM (G). (2 control participants and 2 treated participants were missing data for immunoglobulin concentrations). This analysis included 24 control and 24 rituximab-treated participants.

Among participants being treated with rituximab, IgA levels were highest in participants most recently treated, declining sharply during the first 100 days post-treatment, with no substantial recovery thereafter (range of days since last treatment 0-516). (Figure 3A). Higher cumulative rituximab dosing (range 0-21 total lifetime doses) was associated with persistently low IgA concentrations, which appeared to plateau in individuals who had received more than 5 doses (Figure 3A). These findings suggest a sustained depletion of vaginal mucosal IgA following both recent and repeated rituximab exposure. As expected, peripheral B cells and plasma cells decreased quickly after dosing and remained low. Endocervical B cell proportions demonstrated more variability and relatively less suppression at the same number of doses compared to the periphery, while endocervical plasma cells followed a similar pattern as the periphery (Figure 3B, 3C). Among the participants being treated with rituximab, 14 provided two samples (mean duration between visits was 212 days). Comparison of B-cell and Plasma cell proportions and IgA concentrations among between these two visits did not reveal significant differences in these values within participants. (Supplemental figure 6).

**Figure 3.**
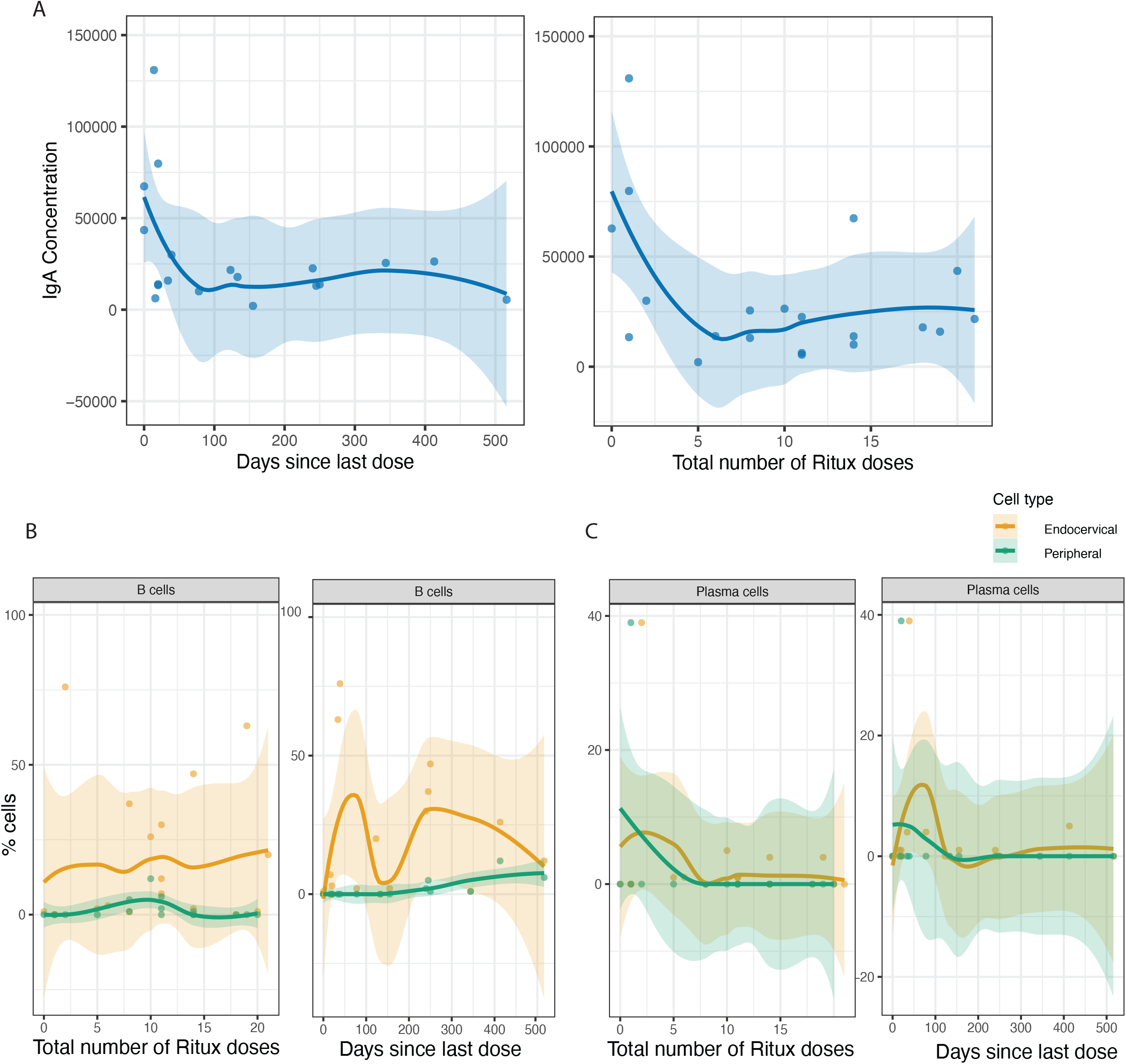
Association between rituximab exposure and genital immune parameters. Vaginal fluid IgA concentrations drop is lower in samples taken a greater number of days since last rituximab dose or after a greater total number of rituximab doses (A). Peripheral B cell proportions are consistently low across samples regardless of time since last dose or total number of doses (B), while endocervical B cells appear to take longer to reach the same low proportions. Plasma cells show a greater similarity between the periphery and the endocervix in association between cellular proportions and duration of treatment or time since treatment (C). Lines represent loess fits with 95% confidence intervals. This analysis included 23 rituximab-treated participants.

To evaluate the relationship between endocervical B cell and plasma cell proportions and vulvovaginal symptoms, we correlated endocervical B cell and plasma cell proportions with a composite symptom score. Across all participants we observed an inverse relationship between symptom score and endocervical B cell and plasma cell proportions, (Figure 4A, 4B), though this association was not statistically significant. Vaginal fluid IgA concentrations (Figure 4C) were not correlated with symptom burden. While these findings are based on a small sample size and are not statistically significant, they align with our prior case series observations, in which most women with inflammatory vaginitis reported resolution of symptoms following discontinuation of rituximab and systemic B cell reconstitution.

**Figure 4.**
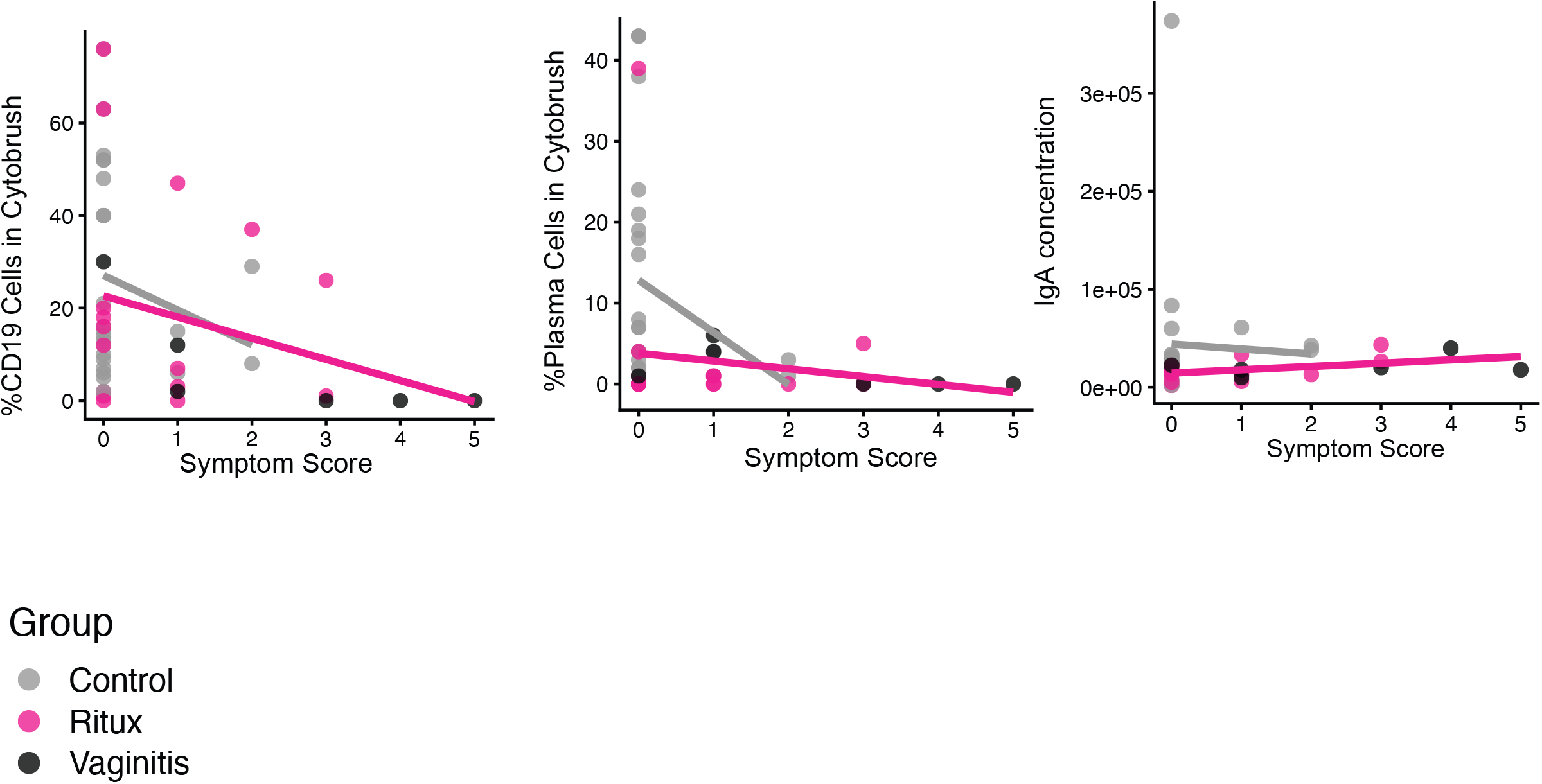
Genital immune parameters and vulvovaginal symptoms. The proportion of endocervical B cells (A) are inversely correlated with the number of moderate-severe vulvovaginal symptoms in control participants, rituximab-treated participants and participants with inflammatory vaginitis. Plasma cell proportions were more strongly associated with symptoms in the control participants (B). Vaginal fluid IgA concentrations were not associated with symptom burden (C). Each point represents one participant, color coded by diagnosis group (rituximab/control/vaginitis). This analysis included 25 control and 25 rituximab-treated participants.

## Discussion

B cells are an understudied component of the cervicovaginal mucosal immune response, and little is known about their role in defense of or homeostasis in the female genital tract. Reports of mucosal inflammatory complications in some individuals receiving B cell–depleting therapies underscore the need to understand whether their mucosal effects parallel or diverge from their well-described systemic effects. In this cohort study, we compared women receiving rituximab for autoimmune disease with healthy controls, assessing endocervical immune populations, vaginal immune mediators, and the vaginal microbiome. Rituximab had measurable effects on the cervicovaginal mucosa, including depletion of B cells and plasma cells, decreased T cell activation, enrichment of proinflammatory cytokines, reduced IgA concentrations, and shifts in the vaginal microbiota. While many of these mucosal changes mirrored known systemic effects, important differences emerged that may offer new insight into the role of B cells in the vaginal mucosa.

As expected, rituximab recipients had a marked reduction in peripheral B cells and plasma cells compared with controls. However, in our study endocervical B cells and plasma cells were reduced to a lesser extent than in the circulation, suggesting that some mucosal B cell populations are distinct and may be relatively protected from systemic depletion. At least one study in mice suggested that B-cell responses to vaginal pathogen challenge were due to circulating B-cells that migrated to the mucosal surface. However our results suggest there may be tissue-resident populations or recruitment from local precursors^14^.Studies in non-human primates have shown mucosal depletion of B-cells and loss of intestinal germinal centers after administration of rituximab,^15–17^ but our study is the first to evaluate these effects in the human uterine cervix, where germinal centers are rare and the biology of local B cells remains largely unexplored.

A primary role of B cells is production of immunoglobulins. Among people being treated with B-cell depleting therapies for autoimmune disorders circulating IgA and IgM are seen to decrease more commonly and more dramatically than IgG.^18–20^ In our cohort, vaginal IgA was significantly lower in the rituximab group, whereas IgM showed no difference. All four subclasses of IgG, which is the most abundant immunoglobulin in vaginal fluid, had comparable concentrations between groups. Some studies show that circulating IgG concentrations decline slowly over prolonged treatment with rituximab,^19,21–23^which we did not observe in endocervical samples. We did not have paired serum measurements to directly compare systemic and vaginal immunoglobulin profiles, but our findings raise the possibility that mucosal IgG may be maintained even when systemic levels decline.

The decline in vaginal IgA is particularly relevant for understanding how rituximab therapy may influence vaginal microbiota, or response to microbiota. In the gut, it is well established that antibodies, primarily IgA, bind microbes and modulate interactions with epithelial cells, mucosal invasion, and expression of inflammatory epitopes^24–27^. In the vagina, IgA-binding of bacteria is more common in *Lactobacillus crispatus* dominant vs. *Lactobacillus* deficient vaginal bacterial communities, suggesting a potential role in maintaining protective microbial composition^28–30^. The reduction in IgA seen in our study implies a smaller available pool for microbial binding, which could influence both microbiome stability and inflammatory response.

Interestingly, despite the higher levels of proinflammatory cytokines such as IL-17, IL-21, IL-23, TNFα, and ITAC in vaginal fluid from rituximab recipients, we found lower activation of endocervical T cells compared with controls. This mirrors observations of reduced circulating activated T cells following rituximab,^31,32^ likely reflecting the disruption of B cell–T cell crosstalk. One possible explanation is that with both B cell and T cell–mediated adaptive immunity suppressed, the vaginal mucosa may rely more heavily on innate immune mechanisms to maintain barrier defense. Heightened activity of epithelial cells, resident innate lymphoid cells, macrophages, or neutrophils could contribute to the observed cytokine enrichment. Some of the cytokines elevated in our cohort, such as IL-17 and IL-23, are also produced by innate immune cells, including γδ T cells and ILC3,^33,34^ and could reflect a compensatory proinflammatory state in response to perceived microbial or tissue signals. This shift toward innate-driven inflammation could help explain why cytokine levels are elevated even as T cell activation is reduced and may indicate that B cells in the FGT normally contribute to dampening or regulating the innate inflammatory responses through antibody-mediated microbial control and immune crosstalk.

Our study has several notable strengths. By leveraging a well-characterized clinical population receiving rituximab for autoimmune disease, we were able to investigate the effects of B cell depletion on mucosal immunity and the vaginal microbiome in a real-world therapeutic context. The limitations of our study include its cross-sectional design that precludes conclusions about causality or the trajectory of mucosal immune recovery over time. Although our longitudinal subset offered some insight into within-participant stability, larger prospective studies will be needed to characterize long-term changes. Additionally, our control group consisted of healthy individuals without autoimmune disease, which limits our ability to isolate the effects of rituximab from those of the underlying autoimmune condition itself. Lastly, while the sample size was sufficient to detect significant differences in multiple immune and microbial parameters, it may not fully capture the variability across different autoimmune diagnoses or treatment regimens.

In this study we found mucosal immune changes in individuals taking rituximab treatment that mirrored in part what is known to happen in the systemic circulation but showed less impact on IgG concentrations and endocervical B-cells than is commonly reported in the circulation. We also found increased mucosal inflammation in people being treated with rituximab. In a small number of participants with inflammatory vaginitis, we showed an inverse correlation between symptom burden and B-cell proportions. Together these findings highlight the unique impact of B-cell depleting therapy on the vaginal mucosa, and its potential for unanticipated adverse side effects.

