## Supplementary figures and images for "An Observational Study of the Impact of Systemic B-cell Depletion on Cervicovaginal Mucosal Environment"

### Supplemental figure 1

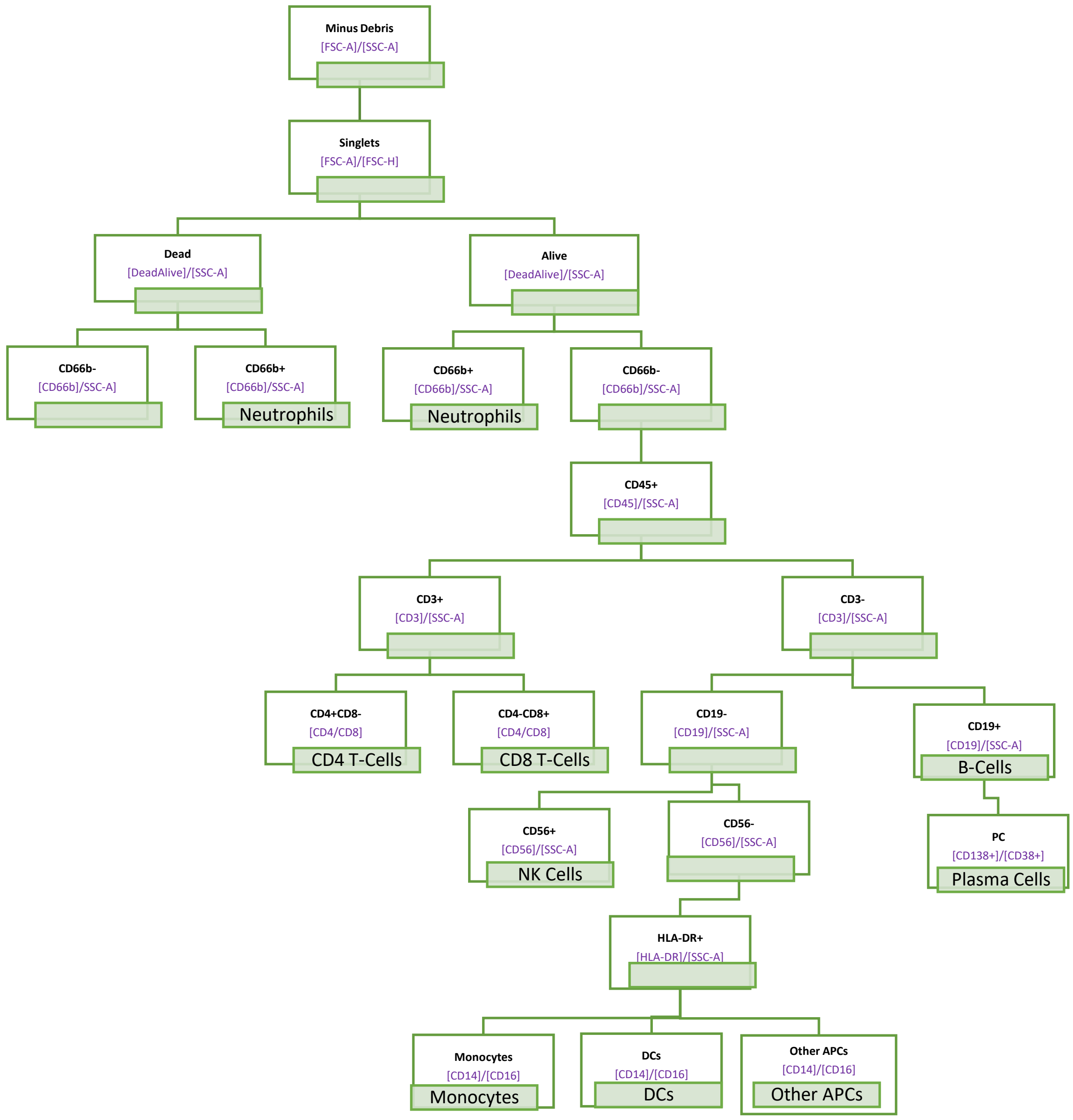

### Supplemental figure 2

Supplemental figure 2

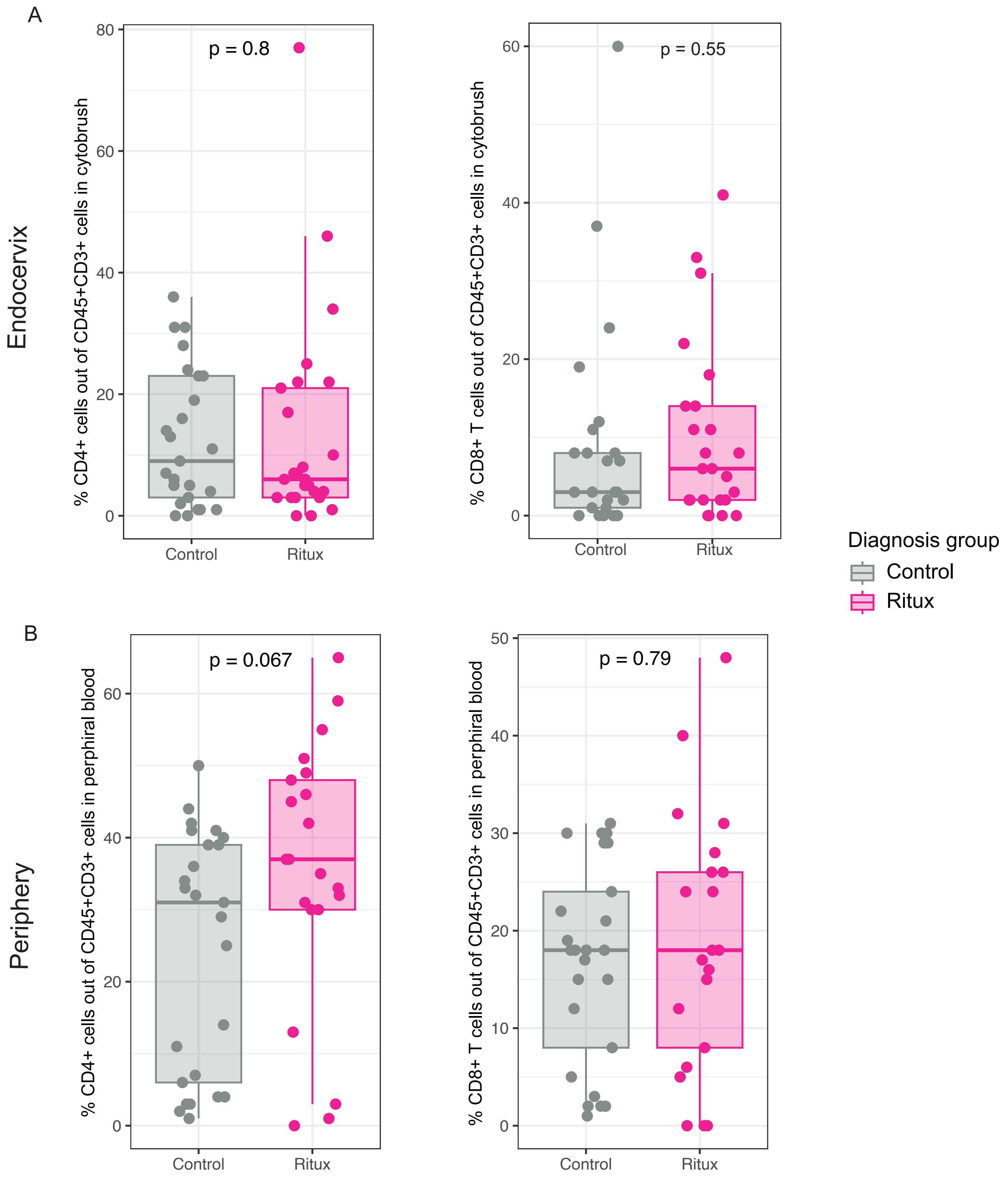

### Supplemental figure 3

Supplemental figure 3

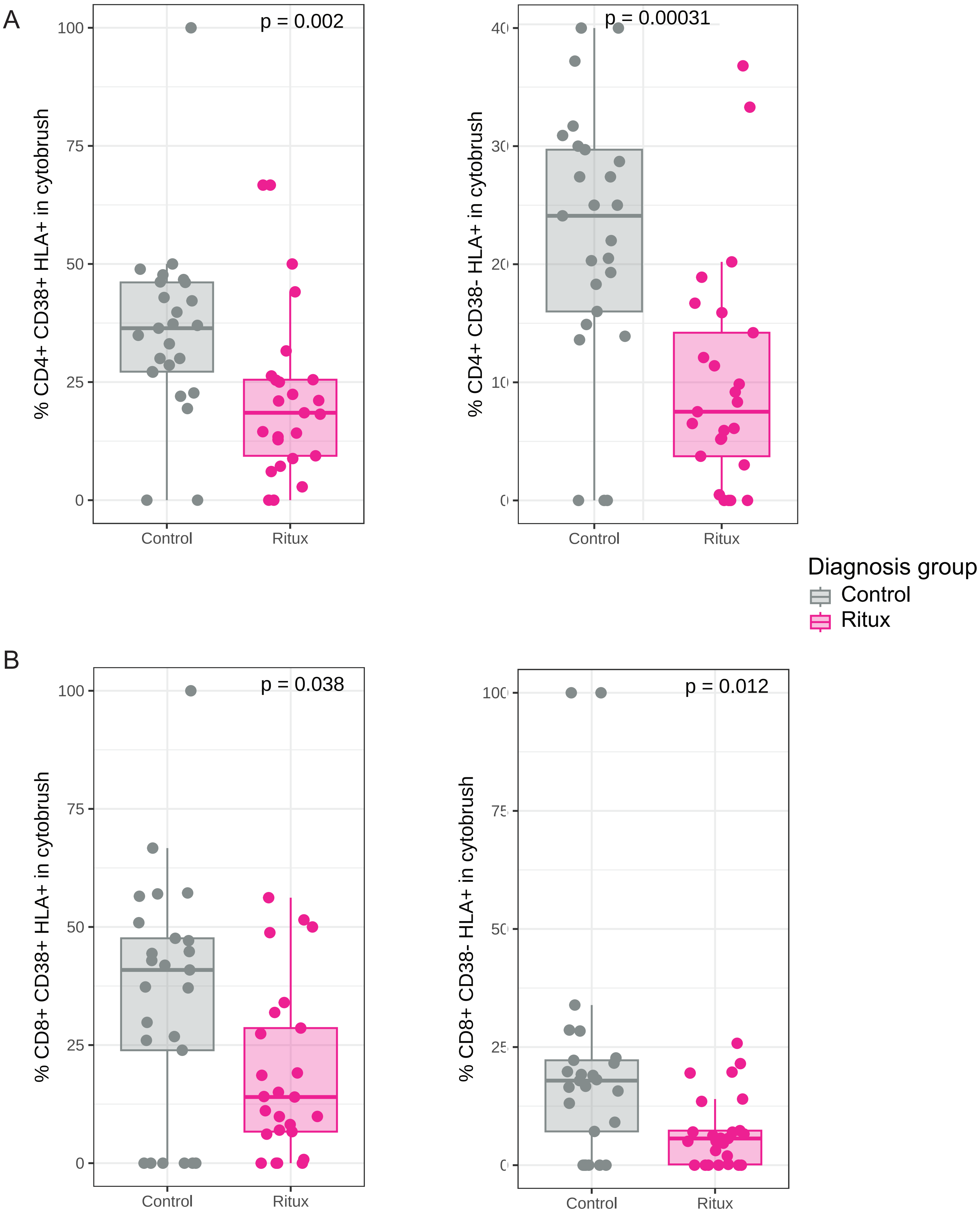

### Supplemental figure 4

Supplemental figure 4

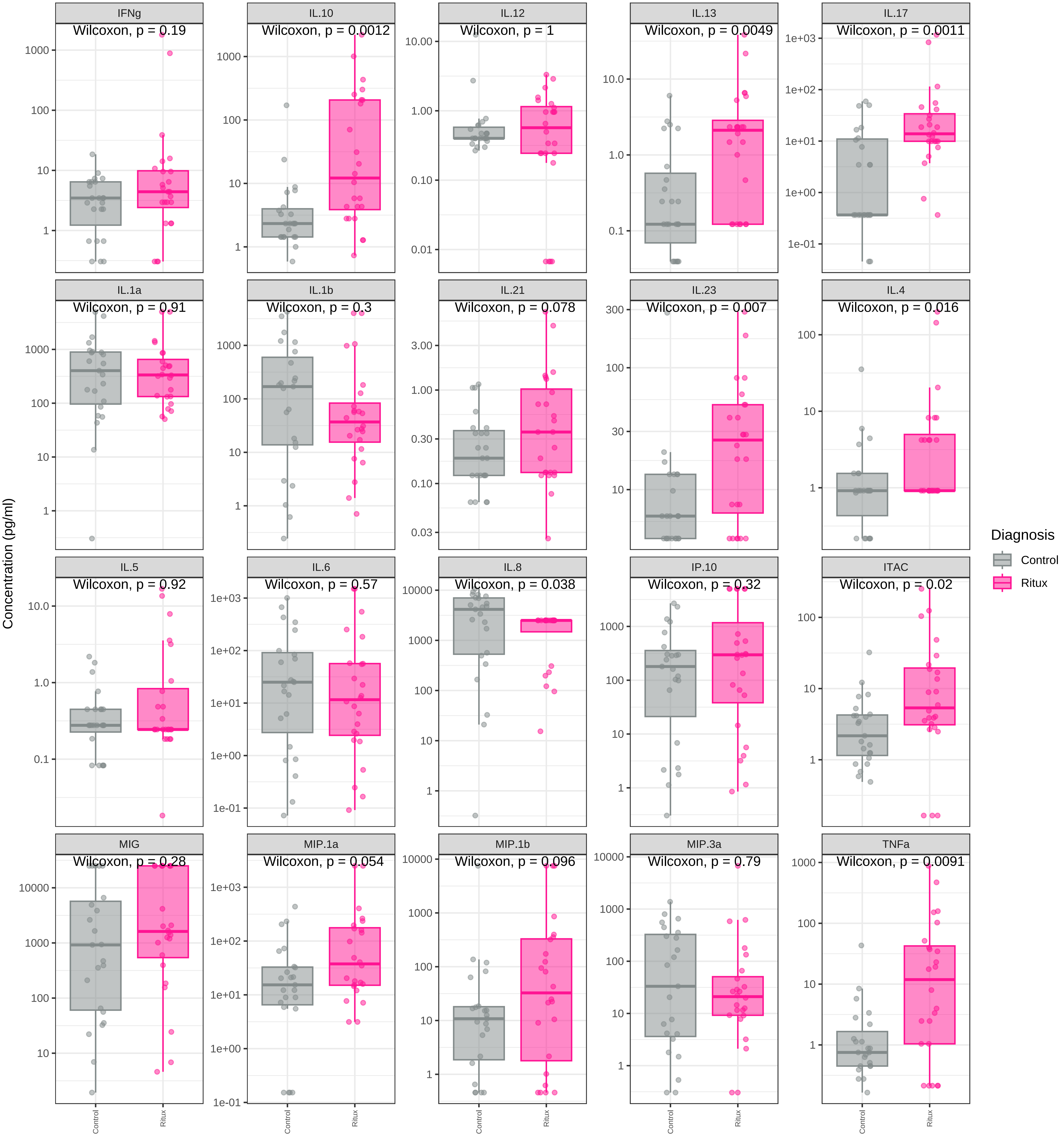

### Supplemental figure 5

#### Supplemental Figure 5

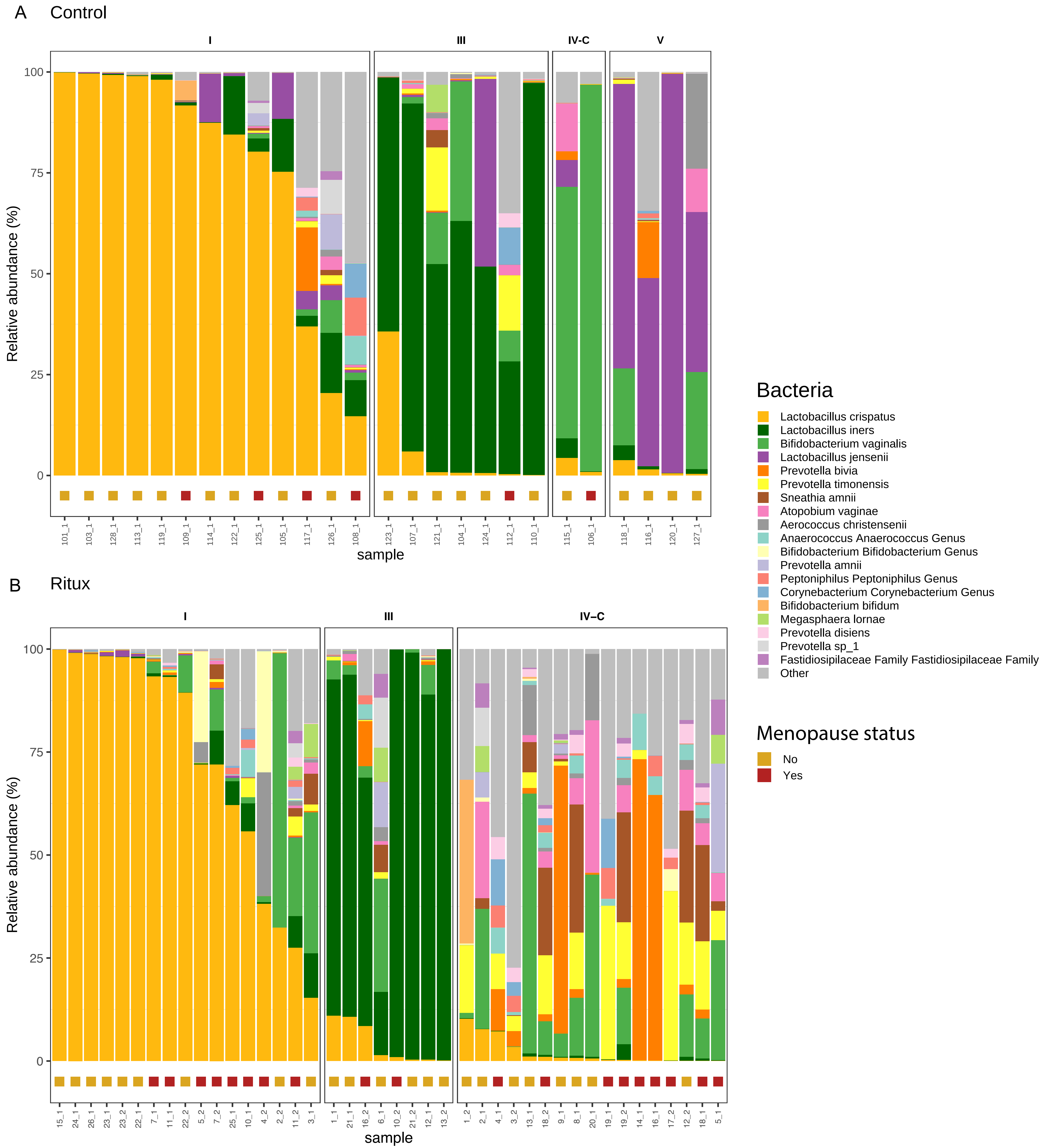

### Supplemental figure 6

#### Supplemental Figure 6

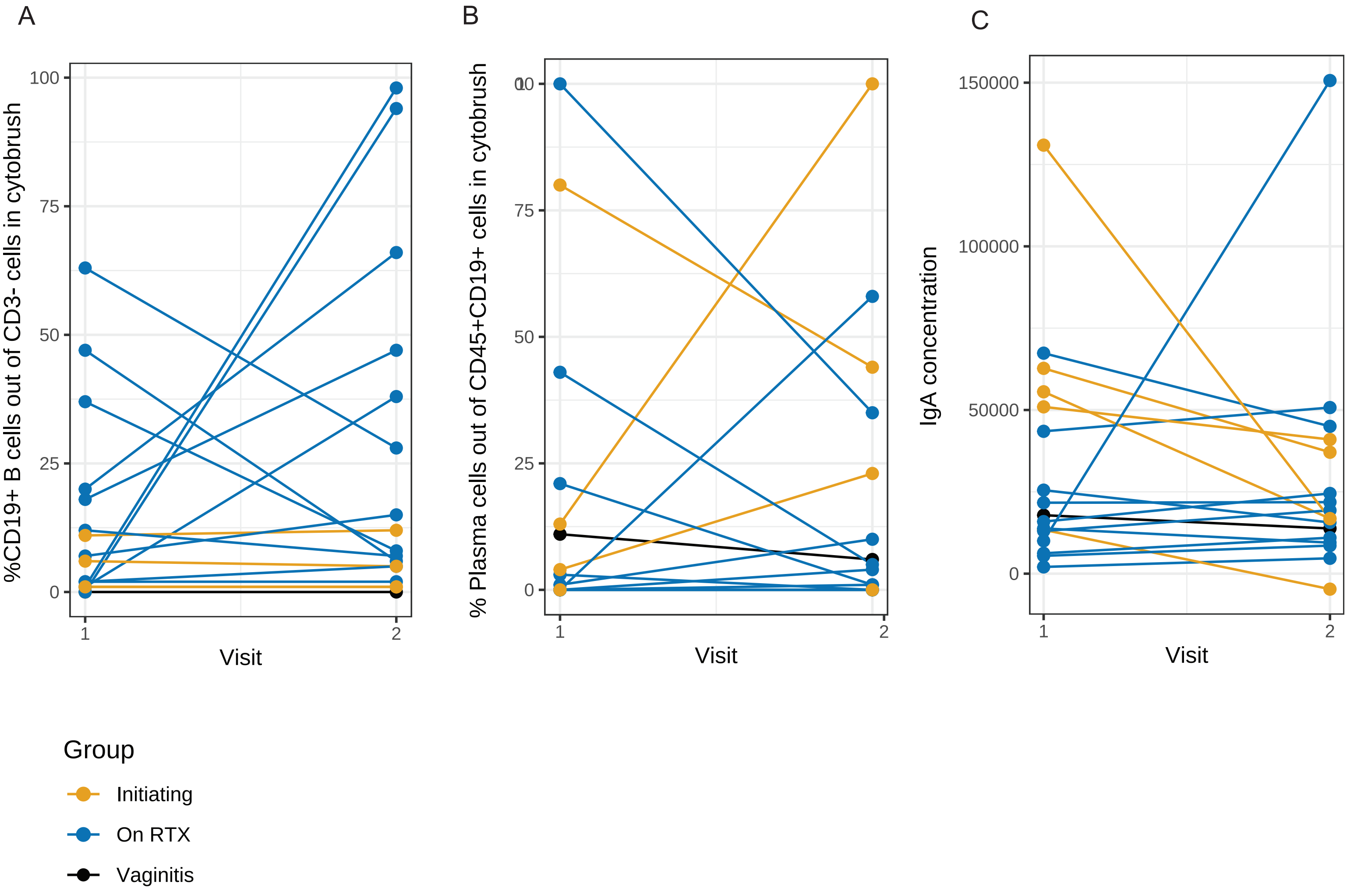
